# Identifying Key Hub Genes that Attribute Varying Host Responses: A Longitudinal RNA-seq Analysis of SARS-CoV-2 Delta and Omicron Infections

**DOI:** 10.1101/2025.06.23.654375

**Authors:** Deewan Bajracharya, Rick J. Jansen, Changhui Yan, Sheela Ramamoorthy

## Abstract

Coronavirus disease 2019 (COVID-19), caused by SARS-CoV-2, has severely impacted global health, with different Variants of Concern (VOCs) resulting in varied clinical outcomes. Among these, the Delta and Omicron variants have drawn significant attention due to their distinct characteristics—Delta is associated with higher virulence and Omicron has greater transmissibility but reduced severity. This study compares host immune responses to the Delta and Omicron variants using RNA-seq data from infected primary human airway epithelial cells. Both variants triggered a robust antiviral innate immune response by 2 days post-infection (dpi). However, Omicron was found to elicit a more rapid immune response, showing pathway enrichment at 1 dpi. In contrast, Delta displayed no immune-related pathway activation within the first 24 hours, suggesting it may evade early immune detection, promoting increased viral replication. By 3 dpi, Delta induced a more aggressive immune response, particularly in pathways related to cell death and pro-inflammatory signaling, such as “programmed cell death” and “regulation of cell death.” Weighted gene co-expression network analysis revealed distinct immune-related genes: Delta infections were characterized by hub genes like MYD88 and IL1R1 involved in pro-inflammatory responses, HLA-A & B, NLRC5 and PSMB9 involved in antigen presentation and TNSF10 and IFR1 involved in pro-apoptotic processes. Conversely, Omicron infections were marked by hub genes such as CXCL1 and CXCL8 involved in immune cells recruitment, MET and LYN involved in reducing hyperinflammatory responses and maintaining immune balance, and IFI44, EIF2AK2 and IFIT5 responsible for the sensing of viral RNA among others.

## Introduction

The coronavirus disease 2019 (COVID-19) pandemic has impacted international communities in an unprecedented magnitude with 777,112,363 cumulative cases and 7,079,587 deaths globally (WHO Coronavirus (COVID-19) Dashboard | WHO Coronavirus (COVID-19) Dashboard With Vaccination Data, n.d.). SARS-CoV-2, the pathogen responsible for COVID- 19, is an RNA virus whose genome codes for genes for the structural proteins (S-glycoprotein, Membrane protein, Envelope protein, and Nucleocapsid protein) and non-structural proteins (components of the Replication Transcription Complex, proteases, helicases, and RNA Dependent RNA Polymerase) (Naqvi et al., 2020). Since being declared the source of a pandemic by WHO in January 2020, the viral genome has undergone several mutations in the aforementioned genomic regions, leading to several variants of SARS-CoV-2.

Although the Wuhan SARS-CoV-2 strain had a slow mutation rate making it unlikely to induce severe health implications (Choi & Smith, 2021), viral infection in immunocompromised hosts has led to an accumulation of multiple mutations, resulting in the emergence of the D614G mutation in the Receptor Binding Domain (RBD) of the spike gene, known for its antibody-neutralizing properties. (Choi & Smith, 2021). Subsequently, among the many SARS-CoV-2 variants that have arisen, a select few have triggered profound global consequences and earned the designation of ’Variants of Concern’ (VOC). These VOC represent distinct lineages of the virus that have elicited global apprehension due to their potential ramifications for public health and disease transmission.

Of all the VOC, two have garnered particular attention due to their unprecedented clinical implications as of August 2023: Delta and Omicron. Delta assumed dominance in multiple global regions, manifesting profound and alarming impacts by exhibiting substantially increased transmissibility compared to preceding variants and increased COVID-19-induced hospitalizations and fatalities. Delta exhibited a twofold increase in the likelihood of hospitalizations, a fourfold increase in the likelihood of hospital admissions, and a 2.5-fold increase in the likelihood of fatalities compared to preceding variants (Choi & Smith, 2021). On the contrary, Omicron is a highly transmissible but less virulent variant (Choi & Smith, 2021). Omicron carries an assemblage of mutations that confer heightened transmissibility compared to previous variants. In a comparative analysis with Delta, which was documented to account for 50% and 80% of total daily infections 100 days after their respective outbreaks, Omicron demonstrated an even more accelerated pace, achieving a higher proportion within just 25 days of its emergence (He et al., 2021). Research findings indicate that Omicron is associated with a reduced likelihood of causing severe disease compared to Delta. Specifically, Omicron is linked to approximately half the rate of hospitalization (fold reduction of 0.6 compared to 1.4) and a roughly tenfold lower mortality rate (fold reduction ranging from 0.03 to 0.3) when compared to Delta (Brüssow, 2022; Ulloa et al., 2022).

Therefore, understanding the molecular mechanisms underlying the disparity in disease outcomes across these VOCs is crucial for developing and maintaining effective diagnostic strategies and vaccines. The majority of existing studies have focused on the differences in host immune response to SARS-CoV-2 in a disease severity-based approach. This study takes a different approach to identify key hub host genes regulating differential host immune responses between the two SARS-CoV-2 VOCs Delta and Omicron. (Figure 1).

**Figure 1.**
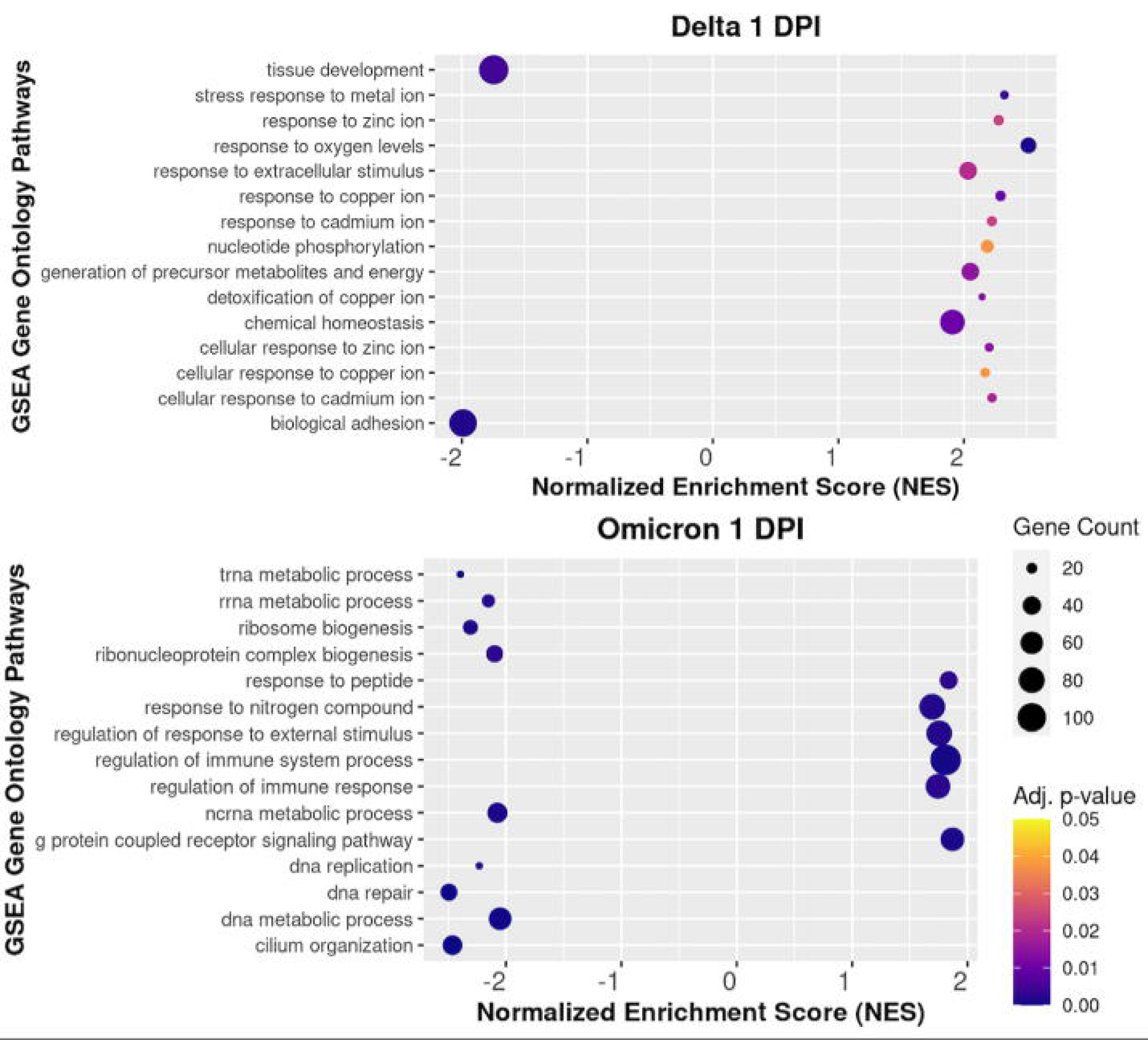
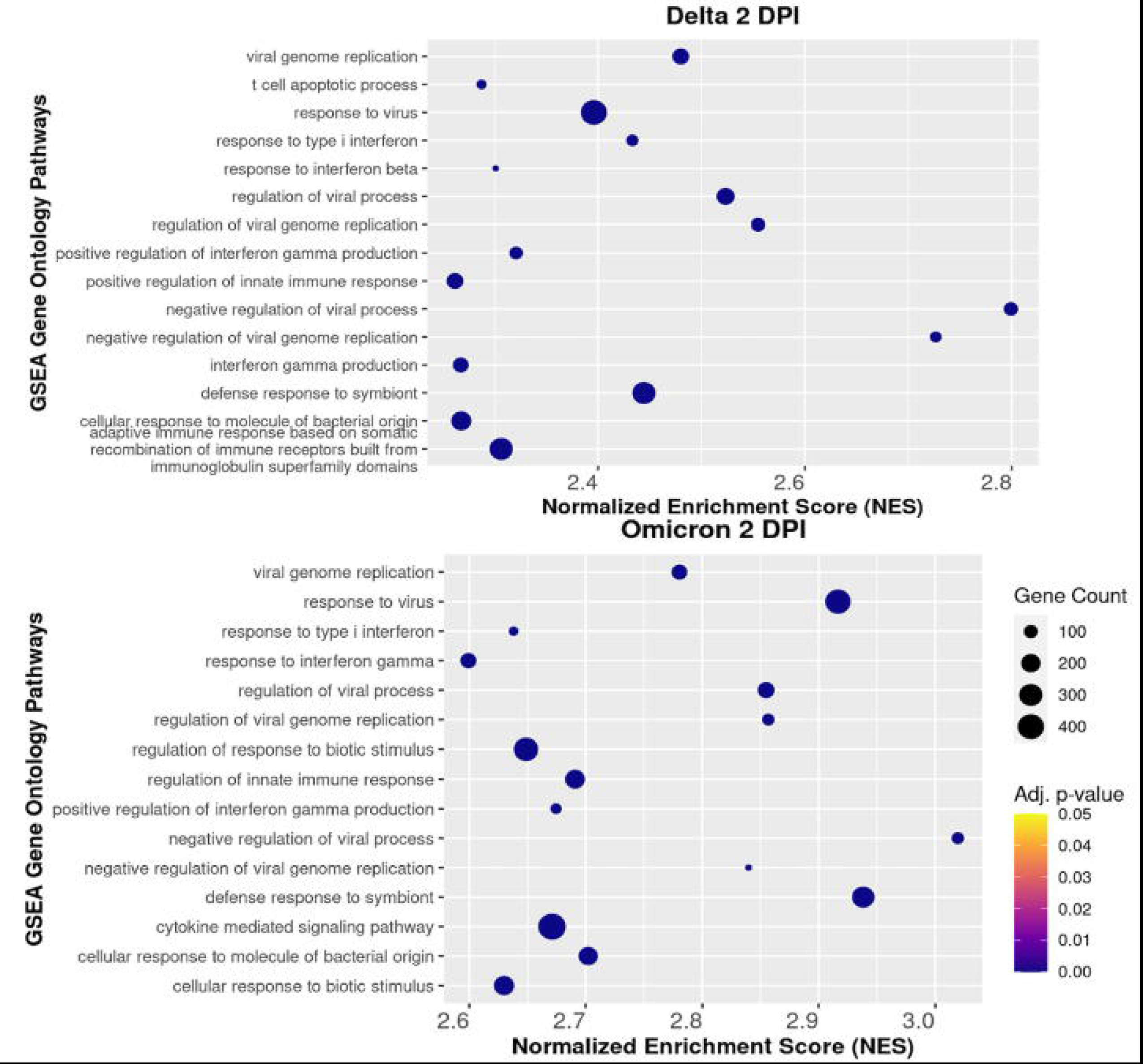
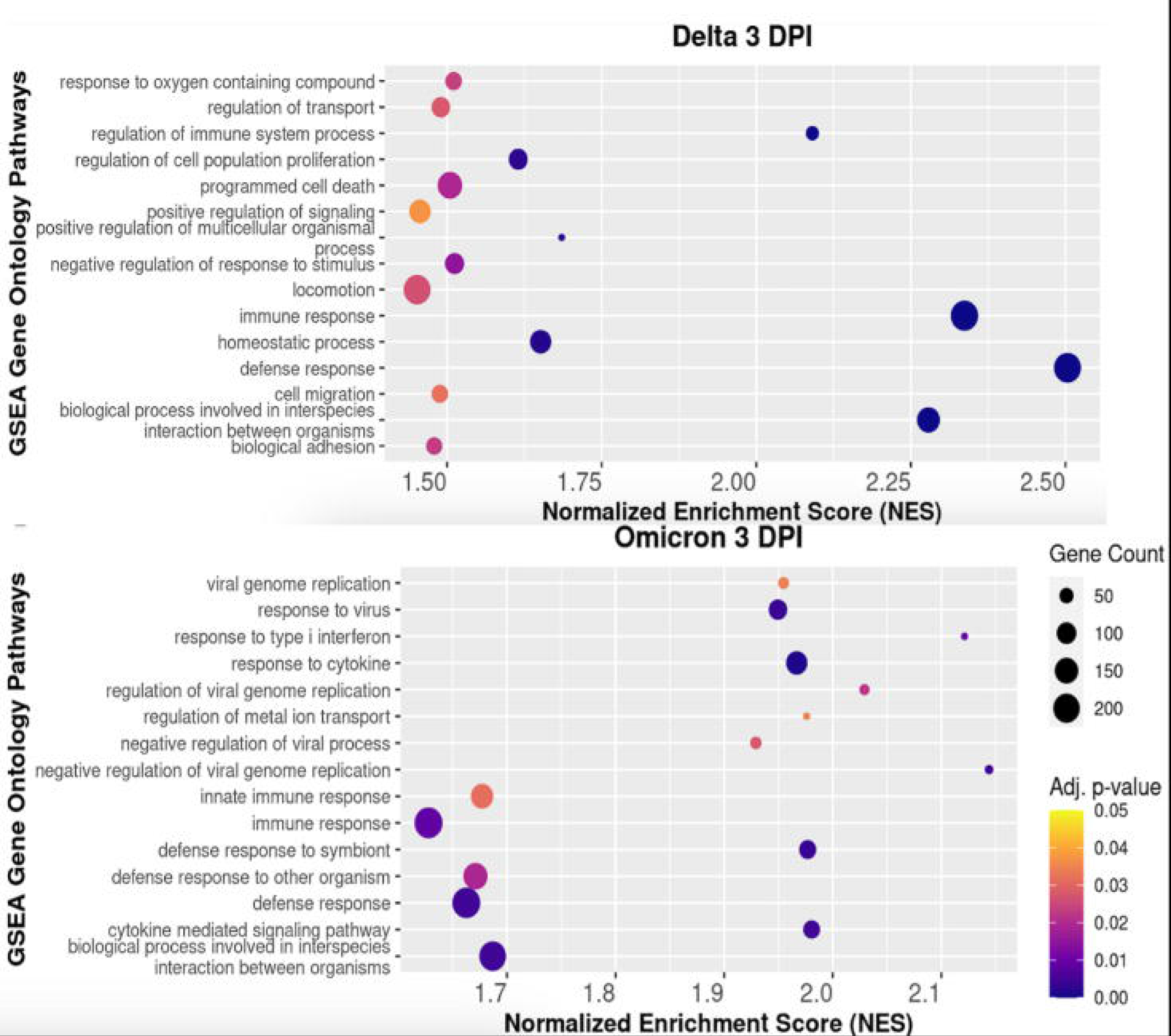
Diagrammatic representation of RNA-seq analysis pipeline from SRR download to identification of hub genes.

## Materials and Methods

### Study Design and Data Acquisition

RNA sequencing (RNA-seq) data for this study were obtained from the Sequence Read Archive (SRA) database at the National Center for Biotechnology Information (NCBI)(Sayers et al., 2022). The gene expression dataset used for this study belongs to the Gene Expression Omnibus (GEO) Accession ID GSE225601 and comprises primary human airway epithelial cells air-liquid interface (ALI) cultures obtained from healthy donors, later infected with Delta (B.1.617.2) and Omicron (B.1.1.529, BA.1). The method of viral inoculation was conserved across all replicates and variants at a final concentration of 105 PFU (plaque forming units). Bulk RNA-seq was performed on ALI cultures infected with respective SARS-CoV-2 variants at 1, 2 and 3 dpi (days post-infection). Mock-infected controls were used as negative controls.

SRA Toolkit was utilized to download the RNA-seq runs corresponding to the Sequence Read Run (SRR) IDs of interest. This tool was chosen for its capabilities in high-throughput data transfer which significantly reduces download times compared to web interfaces as it utilizes multiple threads to download data in parallel.

### Nextflow Pipeline

An automated pipeline has been developed by others using Nextflow (Di Tommaso et al., 2017). This pipeline processes FASTQ files obtained from the NCBI SRA database. The pipeline performs quality control, alignment, mapping and read quantification of each sample in parallel. It outputs a raw gene count matrix for host genes, utilizing a series of established bioinformatics tools for each process as follows:

### Quality Control

Trimmomatic (Bolger et al., 2014) was employed in the preprocessing of the downloaded RNA-seq data to eliminate adapter sequences and remove low-quality bases (Phred score < 20) from the sequencing reads. This process is critical for improving the overall quality and reliability of the data before downstream analyses.

### Mapping Reads to Host Genome

HISAT2 (Kim et al., 2019) was used to align trimmed reads against human reference genome (UCSChg38) indexes. Aligned reads written out to SAM files were sorted and converted into BAM files using SAMtools (H. Li et al., 2009) to retain only host aligned reads.

### Quantification of Host Reads

BAM file reads were annotated to reference annotation file (UCSChg38) using StringTie (Pertea et al., 2015) to generate individual annotated GTF files containing raw and normalized read counts for each sample. StringTie’s prepDE.py was implemented to generate a raw read count matrix from individual annotated GTF files.

### Differential Gene Expression and Gene Set Enrichment Analysis (GSEA)

The DESeq2 package in R (Love et al., 2014) was used for differential expression analyses to identify genes in each experimental group using a selection criteria of an absolute logarithmic fold change (log2FC) value greater than 2, and adjusted P-value < 0.05. (Note: healthy controls were used as negative control groups; Benjamini-Hochberg (BH) method was used to control FDR).

Gene Set Enrichment Analysis, GSEA (Subramanian et al., 2005), which performs enrichment of a list of ranked genes to identify involved biological pathways based on predefined gene sets, was used to unveil functional enrichment of DEGs in each group using clusterProfiler4.0 package and Molecular Signature Database (MSigDB) (Liberzon et al., 2015; Wu et al., 2021). Enrichment results with adjusted P-value < 0.05 were considered statistically significant.

### Co-expression Analysis

Weighted Gene Co-expression Network Analysis (WGCNA) package in R (Langfelder & Horvath, 2008) was used to perform co-expression analysis of host genes in samples infected with respective variant categories. The purpose of performing a co-expression analysis is to identify the cluster of genes correlated with each variant category with an assumption that genes with similar expression patterns across multiple samples are also functionally associated. The read count matrix was tested for outliers in genes and samples using goodGenes and goodSamples functions from WGCNA package. DESeq2 package’s variance stabilization and normalization functions were used. Additionally, a soft thresholding power of 10 was determined based on the maximum R^2^ and minimum mean connectivity criteria using pickSoftThreshold function in WGCNA to obtain an adjacency matrix. The scale-free adjacency matrix was further converted to a topological overlap measure matrix (TOM) to take both pair-wise correlation and correlation of each gene in a pair with shared neighbors into account, generating a robust connectivity measure. Lastly, hierarchical clustering of genes with the least distances (1-TOM) was performed via WGCNA dynamic tree-cut algorithm, with a minimum module size of 30 and deepSplit value of 1. A distance cut-off of 0.25 was chosen to merge modules with high similarity into larger modules.

Each set of genes with similar expression profiles across samples was categorized into a distinct module. Each module was summarized by its module eigengene (ME) value; ME is the first principal component of gene expression values within each module. The modules with the highest correlation with respective traits were selected for additional downstream analyses. Module Membership (MM) was calculated as the correlation of the ME and gene expression for individual genes within the module. MM measures how strongly each gene correlates with the overall expression pattern of a specific module. Genes with higher MM values are considered central or “hub” genes in that module, as their expression aligns closely with the module’s eigengene, representing the core expression pattern of the module. Gene Significance (GS) values were calculated as the correlation of individual gene’s expression with trait of interest (Mock, Delta, and Omicron). Genes with higher GS values are considered more relevant to the biological trait of interest as their expression is closely related to variations in the trait.

### Hub Genes for Each Variant Category

The genes with highest GS and MM values (|GS| > 0.6 and |MM| > 0.8) for Delta and (|GS| > 0.5 and |MM| > 0.8) for Omicron were then selected for additional downstream identification of candidate hub genes. Identified DEGs for each VOC category that overlapped with module of interest were classified as candidate hub genes.

### Protein-Protein Interaction (PPI) Network for Candidate Hub

Candidate hub genes with highest MM and GS values within each module for respective VOC categories were plugged into Cytoscape (Shannon et al., 2003) to generate respective PPI using STRING database (Szklarczyk et al., 2021). The software cytoHubba (Chin et al., 2014) was used to identify ten key hub genes or central nodes from each PPI network using Maximum Clique Centrality (MCC) algorithm. The MCC algorithm ranks nodes based on network centrality, highlighting genes with the highest interaction potential. Key hub genes represent the most central and influential node within the network.

### Ethical Statement

Since all the data obtained for the purposes of this ecological study were accessed from publicly available and open resources, no IRB or ethical approval was needed.

## Results

### Expression of Host Genes in SARS-CoV-2 Infected ALI Cultures

The read counts for ALI cultures infected with Delta (3 replicates at 1, 2 and 3 dpi), and Omicron (3 replicates at 1, 2 and 3 dpi) were compared with those for healthy controls (3 replicates at 1, 2 and 3 dpi). A total of 2445, 1148 and 1642 DEGs were identified in ALI infected with Delta variant and 1333, 597 and 715 DEGs were identified in ALI infected with Omicron variant at 1, 2 and 3 dpi respectively (Figure S1).

### Functional Annotation of DEGs

GSEA was performed using clusterProfiler4.0 to map the DEG for each variant category against Gene Ontology Biological Process (GO BP) subcategory in MSigDB. Both VOCs induced statistically significant sets of pathway enrichment across all three timepoints.

The patterns of GO terms enriched in the Delta category of host genes at 1 dpi comprise pathways involved in several metal ion responses whereas pathways involved in “tissue development” and “biological adhesion” were inhibited (Figure 2a). There were no immune-related terms enriched by Delta variant category at 1 dpi. The patterns of GO terms enriched by the Omicron variant category at 1 dpi reveal GO terms such as “regulation of immune system” and “regulation of immune response” being upregulated and pathways involved in tRNA, rRNA and ribosome being downregulated (Figure 2b). Absence of immune-related enriched terms at 1 dpi in Delta variant category potentially indicates the ability of Delta variant to evade immune cells during the early phase of infection as well as the well documented higher replication rate of Omicron variant within 24 hpi (Castaneda et al., 2023; Tanneti et al., 2024).

**Figure 2.**
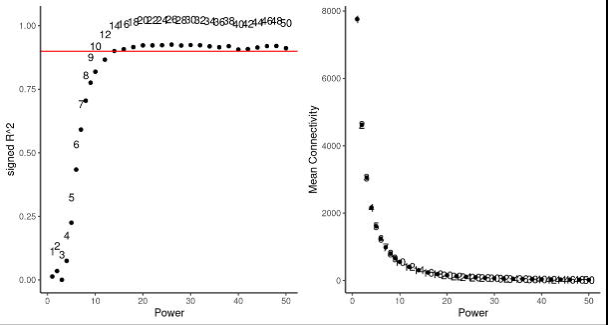
GSEA plot showing the most enriched host pathways in (a) Delta, and (b) Omicron infected hosts at 1 dpi .

At 2 dpi, both categories of host genes were found to enrich several immune response pathways (Figure 3). GO terms like "negative regulation of viral process" and "response to type I interferon" were enriched in both variant categories. This suggests a coordinated cellular response to suppress viral replication and activation of antiviral pathways mediated by type I interferon signaling. While Omicron-infected cells displayed a stronger immune response with enriched pathways predominantly associated with antiviral and immune signaling mechanisms such as “viral genome replication,” “response to type I interferon,” “response to interferon beta,” and “regulation of viral process”, Delta displayed enrichment of “T cell apoptosis”. This is consistent with findings from a study by Saichi et al., 2021, which states that SARS-CoV-2 induces pro-apoptotic pathways in immune cells including T-cells as an immune evasive strategy.

**Figure 3.**
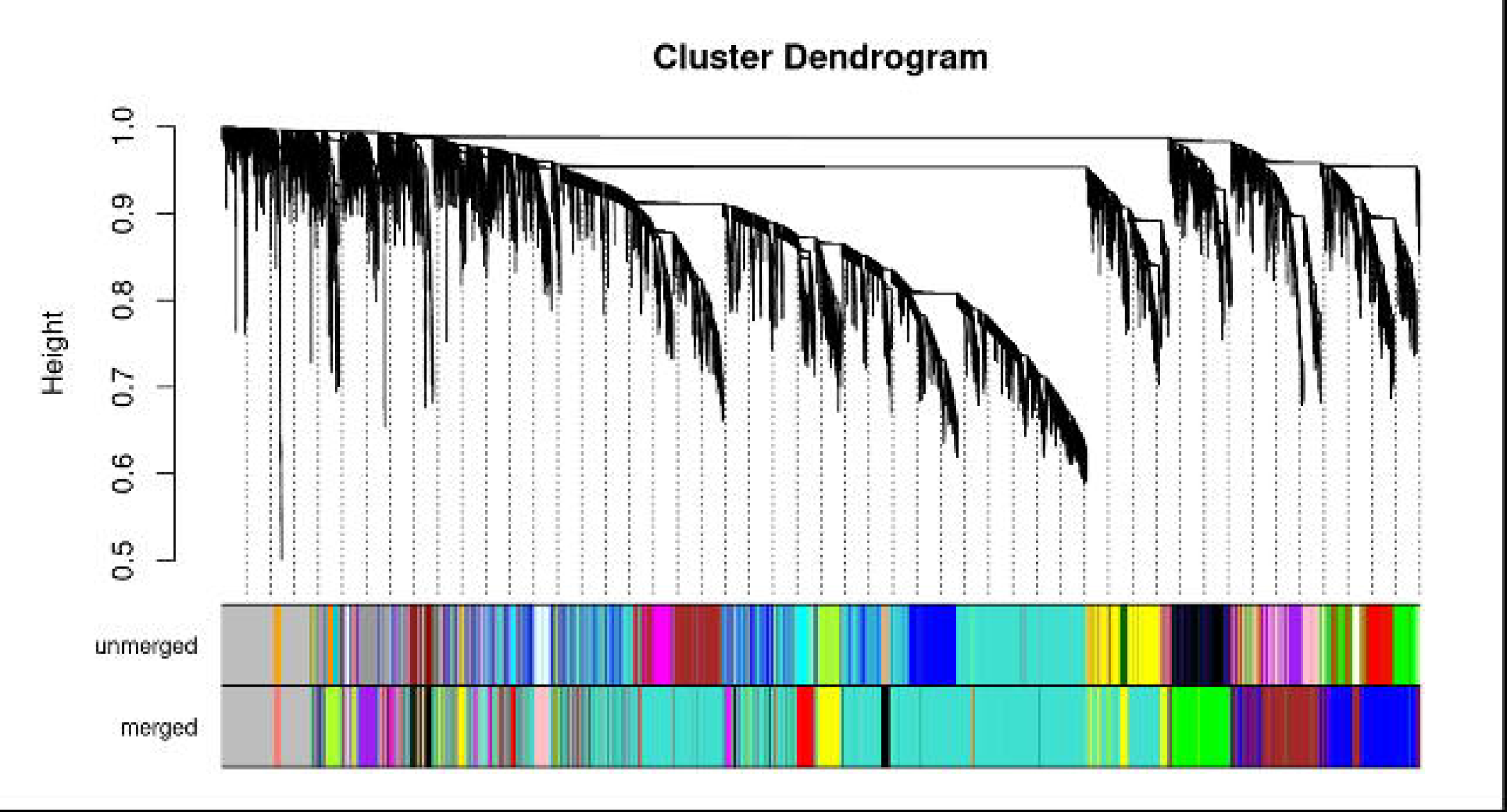
GSEA plot showing the most enriched host pathways in (a) Delta, and (b) Omicron infected hosts at 2 dpi.

At 3 dpi, Delta variant category showed an aggressive immune response with enrichment of GO term" programmed cell death", "regulation of cell death" (Figure 4a). This indicates a heightened antiviral response with the involvement of apoptotic pathways. Along with the previously observed enrichments in "response to type I interferon", "innate immune response" and "negative regulation of viral processes" indicating a sustained antiviral response at 3 dpi, Omicron variant category also displayed enrichment of "Cytokine mediated signaling pathways" and "Response to Cytokine" (Figure 4b). This suggests potential recruitment and activation of immune cells through cytokine signaling pathways. The continued presence of terms related to interferon signaling ("response to type I interferon" and "response to interferon beta") further emphasizes the ongoing antiviral efforts.

**Figure 4.**
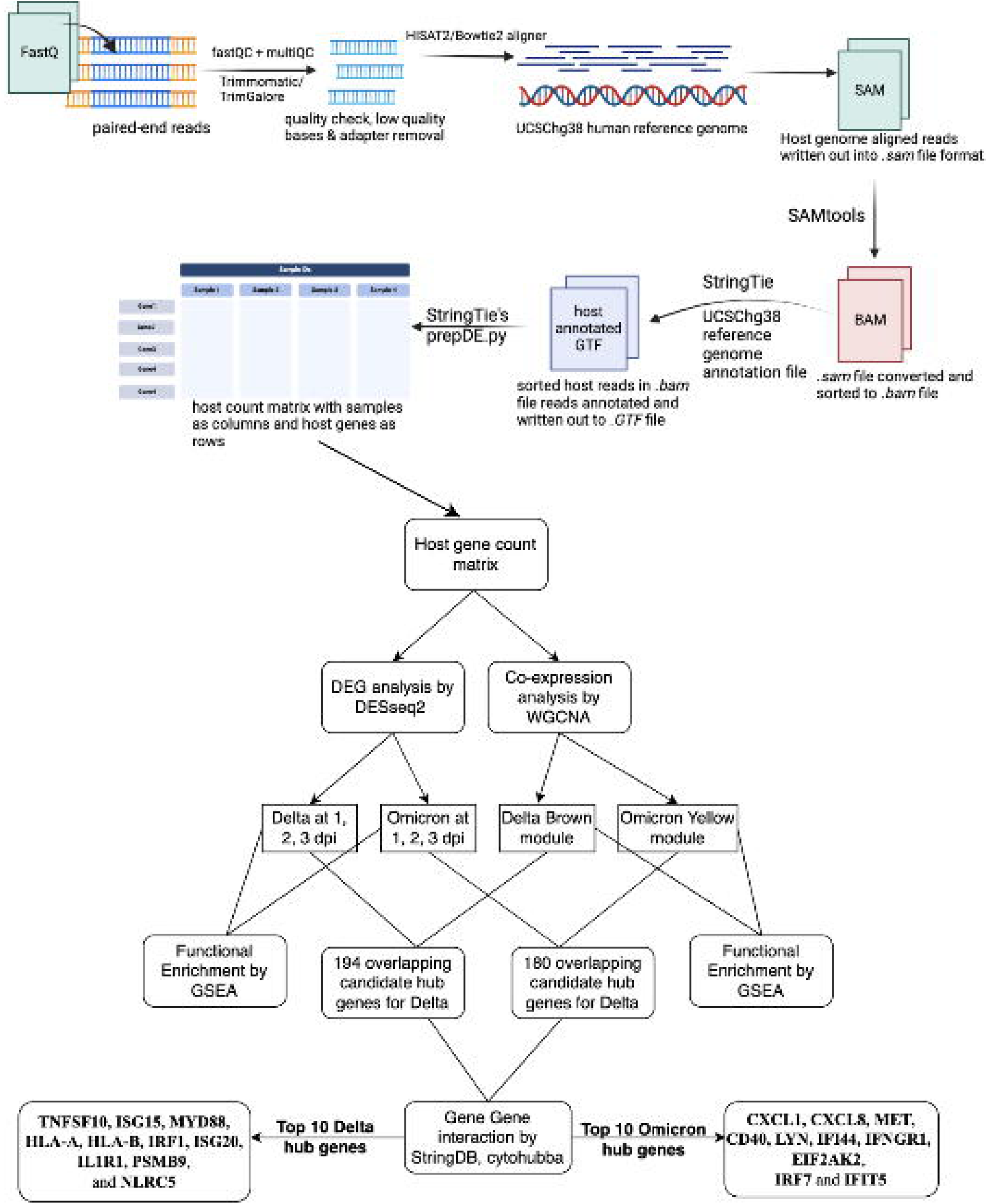
GSEA plot showing the most enriched host pathways in (a) Delta, and (b) Omicron infected hosts at 3 dpi.

### Identification of WGCNA Modules and Module-Trait Association

WGCNA was performed on the read count matrix to identify sets of host genes, referred to as modules, for each variant category. These modules comprise genes whose expression profiles exhibit the most statistically significant correlation with the presence of the respective variant category. The analysis was conducted using a soft-threshold β of 10, as shown in Figure 5a. Dynamic tree cut was implemented to generate a total of 14 co-expression modules, each module consisting of a unique set of host genes with correlated expression profiles as shown in Figure 5b.

**Figure 5.**
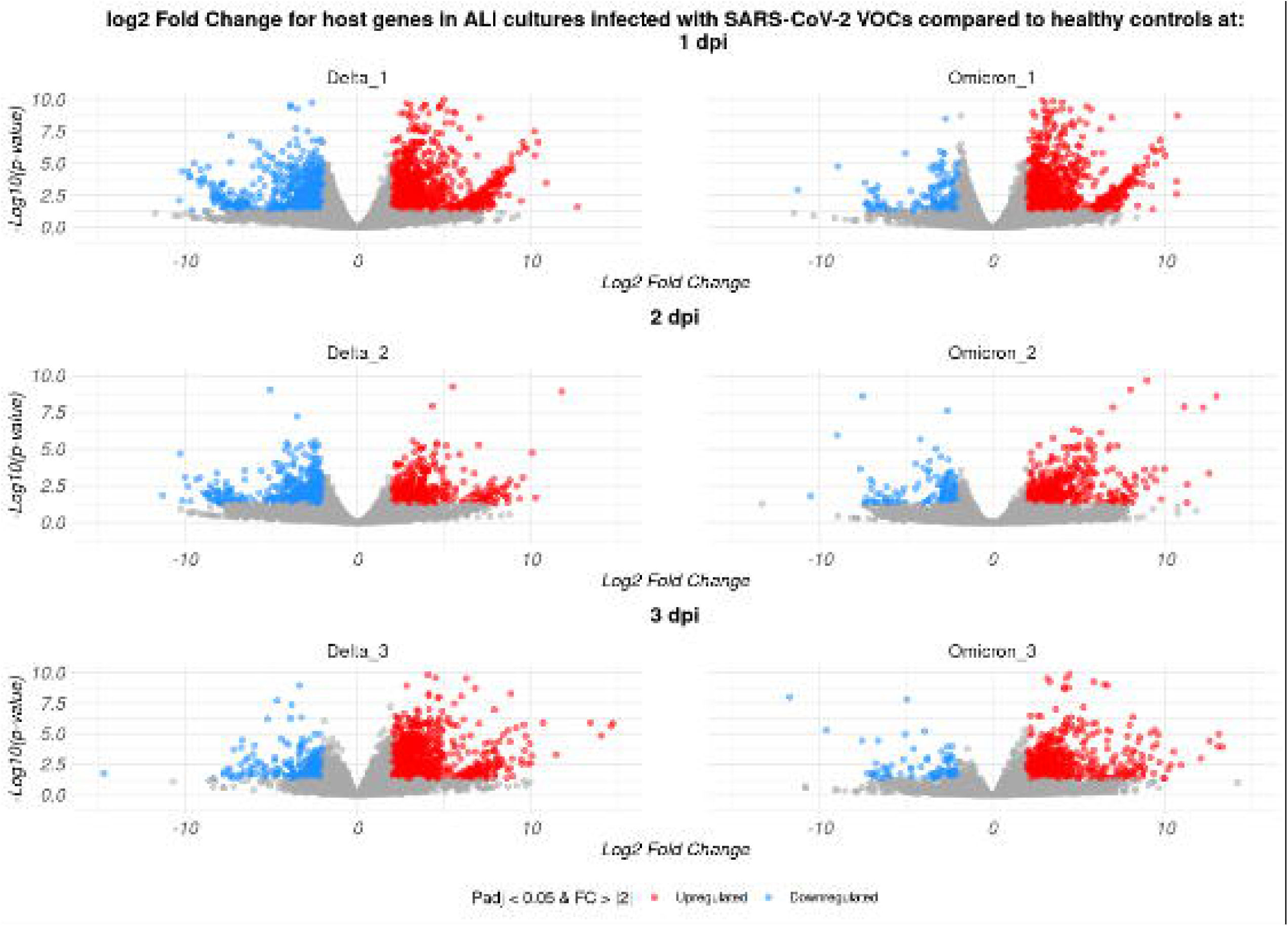
Identification of key modules correlated with Delta and Omicron infections using WGCNA. (a) Analysis of the scale-free topology fit index (left) and the mean connectivity (right) for various soft-thresholding powers. (b) Clustering dendrograms of genes based on a dissimilarity measure (1-TOM).

Module-trait association analysis was performed between ME for each module and outer features (presence of respective variant categories or mock) and visualized by heatmap profiles as shown in Figure 6. The results confirmed the brown module to be the most correlated with Delta variant infections (r = 0.57, p-value < 0.001) and negatively correlated with the mock trait category (r = -0.85, p-value < 0.001). Similarly, the yellow module was found to be the most correlated with Omicron variant infections (r = 0.53, p-value < 0.001) and negatively correlated with mock trait category (r = -0.31, p-value < 0.05). These findings imply that the genes comprising brown and yellow modules were upregulated in the presence of respective VOCs. As shown in Figure S2a, MM in the brown module positively correlated with GS for Delta infection (correlation = 0.36, p-value < 0.001). This indicates that the genes that are most significantly associated with Delta infection were found to share modality within the brown module.

**Figure 6.**
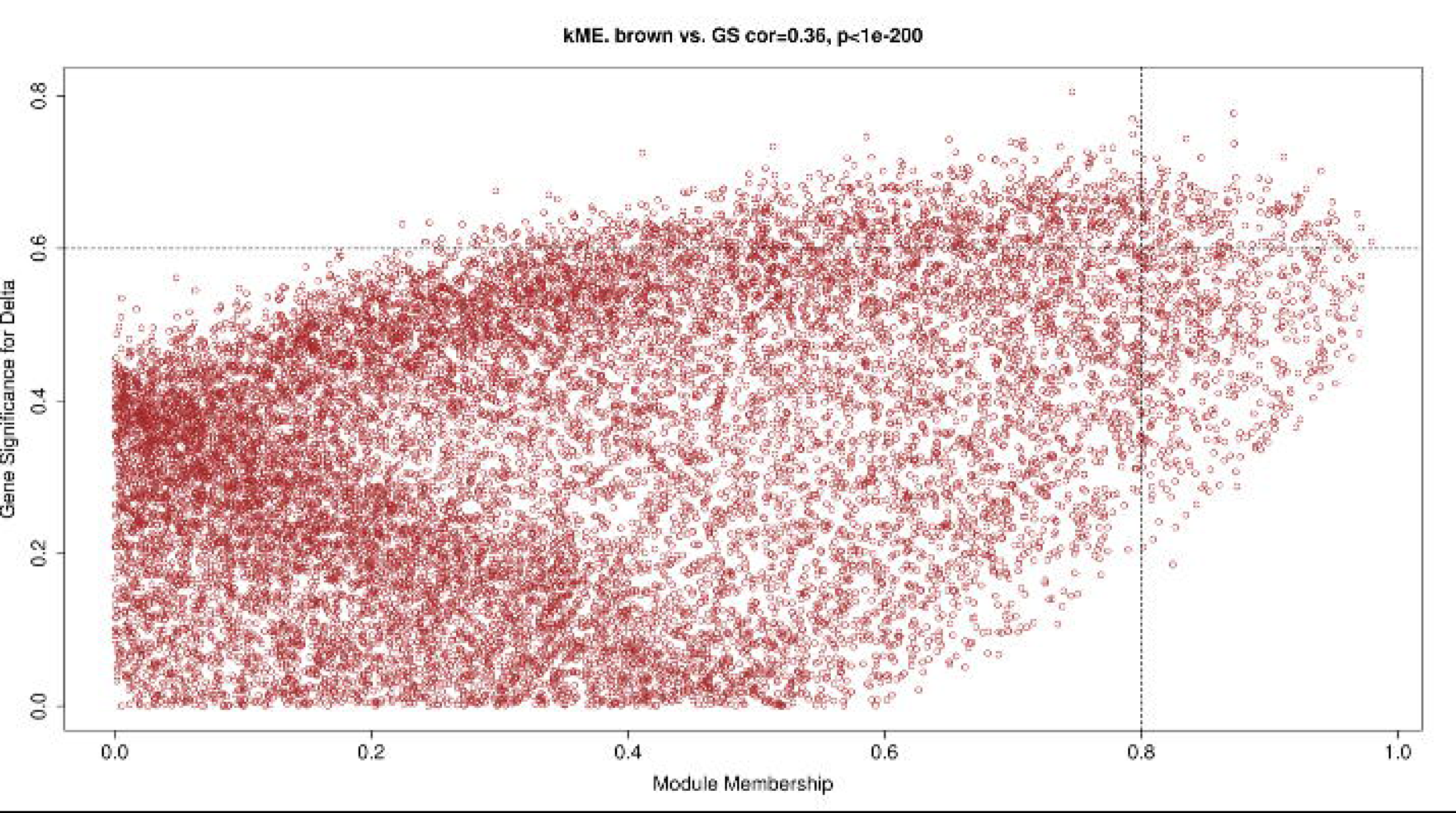
Identification of key modules correlated with Delta and Omicron infections using WGCNA’s Module-trait associations between module eigengenes and sample traits. Each cell contains the correlation coefficient and P value. Note: *p < 0.05; **p < 0.01; ***p < 0.001.

Likewise, as shown in Figure S2b, MM in the yellow module positively correlated with GS for Omicron infection (correlation = 0.42, p-value < 0.001), suggesting that genes that are most significantly associated with Omicron infection were also the key genes in the yellow module.

### Candidate Hub Gene Identification for Modules of Interest

As shown in Figures S2a and S2b, genes in the top left corner of the MM-GS plot within the brown and yellow modules were identified as the genes most associated with respective VOC infection and hence can be termed candidate hub genes. A total of 1168 and 1026 genes were selected as co-expressed genes within the brown and yellow modules respectively. Additionally, the DEGs for each VOC category were compared to the candidate genes, and their overlaps were selected for further screening with the goal of identifying the top candidate hub genes, resulting in a total of 194 and 180 candidate hub genes within the brown and yellow modules respectively.

### Identification and Evaluation of Top Hub Genes between Delta and Omicron

194 and 180 candidate genes for the Delta and Omicron VOC categories respectively were uploaded to Cytoscape to create a PPI using STRING database. A PPI with 184 nodes and 1207 edges that displayed protein interactions involved with the list of gene names in Delta VOC category was constructed. Similarly, a PPI with 158 and 437 edges that displayed protein interactions involved with the list of gene names in Omicron VOC category was constructed.

cytoHubba was implemented onto each PPI network to identify the top 10 central nodes ranked by Maximal Clique Centrality (MCC) algorithm, which were termed as key hub genes for each module of genes (brown and yellow) as shown in Figure S3. TNFSF10, ISG15, MYD88, HLA-A, HLA-B, IRF1, ISG20, IL1R1, PSMB9, and NLRC5 were identified as the 10 hub genes for the Delta VOC category. CXCL1, CXCL8, MET, CD40, LYN, IFI44, IFNGR1, EIF2AK2, IRF7 and IFIT5 were identified as the 10 hub genes for the Omicron VOC category. These 10 hub genes had the most influential roles in the host response to the respective VOC category. The MCC algorithm in cytoHubba ranks each node in a PPI network based on the maximum number of cliques, which are groups of nodes where every node is connected to every other node. This ranking reflects the weighted centrality of each node, based on the size of all the cliques it represents, thereby quantifying the interactions of each node within the network to identify the most central and potentially influential hub nodes (Chin et al., 2014).

## Discussion

Delta and Omicron variants have both demonstrated varying disease manifestations during the COVID-19 outbreak with the former resulting in more virulent consequences and the latter displaying higher transmissibility (Choi & Smith, 2021; Goethem et al., 2022). While there have been several studies attributing host immune responses to disease severity (Girona-Alarcon et al., 2022; Karami et al., 2021; Ulloa et al., 2022), evaluation of gene expression profiles differences between Delta and Omicron variants to identify key hub genes attributing to disease manifestations needs to be further investigated. Our study implements a combination of DEG analysis and WGCNA on RNA-seq datasets obtained from primary human airway epithelial cells air-liquid interface (ALI) cultures infected with SARS-CoV-2 variants. This process is used to identify the key hub genes and consequently the pathways they regulate that are distinct to each VOC category. This cell type was selected because of its close resemblance with the human airway epithelia, where the virus primarily infects (Castaneda et al., 2023). The Delta and Omicron variants were identified as the major VOCs for this study due to their association with more severe disease outcomes despite US vaccination efforts as we reported in a previous study (Bajracharya & Jansen, 2024), highlighting the need for proper investigation of the molecular mechanisms that differ in hosts challenged with these two primary variants.

A viral infection is capable of stimulating a cascade of signaling pathways within an infected host cell, resulting in variation in the host transcriptome to stimulate essential immune responses against the viral agent (Thaker et al., 2019). Our study utilized the transcriptome expression datasets and focused on the differences in host immune responses to Delta and Omicron infections. With selection criteria of |log2FoldChange| > 2 and adjusted p-value < 0.05, the GSEA of DEGs revealed that both variants induce a more robust innate immune response in the host culture. While a consistent set of antiviral innate immune GO pathways such as response to type I Interferons, Interferon alpha and beta and ISG that inhibit viral replication were found to be upregulated between both VOCs by 2 dpi, the GSEA did not detect any immune-related pathway enrichment in Delta infections for the first 24 hours after inoculation (1 dpi).

Conversely, Omicron infections induced immune responses within the first 24 hours. These findings may provide evidence to support the previously identified higher replication rate of the Omicron variant compared to Delta in certain cell types, including upper respiratory epithelial cells (Castaneda et al., 2023; Tanneti et al., 2024). This rapid replication may allow Omicron to achieve high viral loads before the innate immune response can fully contain it. Even with an early innate immune response, the sheer rate of Omicron replication may overwhelm the host’s ability to clear the virus quickly, prolonging the infection. Additionally, Omicron has evolved mutations in its spike protein and other viral components that allow it to evade neutralizing antibodies and possibly certain innate immune detection pathways. This could dampen the effectiveness of the early immune response despite its activation. For instance, the early induction of type I interferons and interferon-stimulated genes (ISGs) may not fully counteract Omicron’s replication because of its ability to interfere with antiviral pathways. Omicron’s preference for replication in the upper respiratory tract may result in a different immune landscape compared to Delta, which has been associated with more severe lower respiratory tract infections. This phenomenon may reflect the Omicron’s enhanced ability to replicate efficiently in the upper respiratory tract, leading to early and robust detection by innate immune sensors such as pattern recognition receptors (PRRs) (Hui et al., 2022; McMahan et al., 2022).

Omicron’s increased reliance on cathepsin-mediated entry mechanisms rather than TMPRSS2- dependent pathways may also play a role in its ability to evade certain host barriers and replicate more efficiently in specific cellular environments (Hui et al., 2022). This could further explain the rapid immune activation observed, as viral replication in susceptible upper airway cells might lead to the quicker release of viral RNA and proteins that stimulate immune pathways.

Additionally, the lack of an early immune response in Delta infections may reflect the variant’s ability to evade antiviral pattern recognition receptors (PRRs) detection at low viral loads. This immune evasion could be a critical factor allowing the Delta variant to establish a robust infection before the host immune system is activated. Notably, distinct Gene Ontology (GO) term enrichments observed in Delta-infected air-liquid interface (ALI) cultures indicate the regulation of cell death and apoptotic signaling pathways, alongside other antiviral innate immune response pathways, at the 3 dpi timepoint. These findings align with clinical studies reporting severe cytopathic effects and significant host cell death during Delta variant infections (Choi & Smith, 2021; Lee et al., 2022; Tanneti et al., 2024). This regulation of cell death and apoptosis may serve dual purposes for the Delta variant. On one hand, it could be a result of the virus inducing host cell death to propagate infection. On the other hand, such processes might also represent a host defensive mechanism attempting to limit viral replication through infected cell clearance. Furthermore, the severe cytopathic effects observed in Delta infections are consistent with its enhanced ability to cause epithelial barrier disruption, loss of ciliated cells, and widespread tissue damage compared to other variants (Tanneti et al., 2024). These observations highlight Delta’s aggressive pathogenic mechanisms, further underscoring its capacity to evade early immune responses while inducing significant cellular damage later in the infection process.

Following the DE analysis, gene co-expression analysis using WGCNA provided further insight into the most significantly correlated modules for Delta and Omicron infections. Of the 14 identified modules generated, the brown and yellow modules were found to be the most positively correlated with Delta and Omicron infections respectively while being most negatively correlated with mock infections, suggesting the genes within these modules were upregulated as an immune response to respective VOC infections. Gene set enrichment analysis of the brown module revealed the involvement of pathways critical for antiviral innate immune responses.

These pathways include interferon-alpha (IFN-α) and interferon-beta (IFN-β) signaling, antigen-presenting cell activation, defense response to viruses, and the positive regulation of programmed cell death. These findings suggest that the brown module is primarily associated with the activation of innate immune mechanisms aimed at controlling viral replication and initiating adaptive immune responses. On the other hand, gene set enrichment of the yellow module highlighted pathways associated with broader immune regulatory functions. These pathways include immune regulation, defense response, response to external stimuli, immune effector production, and other general antiviral immune pathways. This indicates that the yellow module encompasses a wider range of immune processes, reflecting its role in orchestrating both innate and adaptive immune responses against viral infections.

The identification of hub genes within each module, using the cytoHubba tool, revealed key molecular driver genes that are highly correlated with the host’s response to Delta and Omicron infections. For Delta infections, the hub genes TNFSF10, ISG15, MYD88, HLA-A, HLA-B, IRF1, ISG20, IL1R1, PSMB9, and NLRC5 were identified as central nodes in the interaction network. These genes are heavily implicated in antiviral responses, immune modulation, and apoptosis regulation. For Omicron infections, the hub genes identified were CXCL1, CXCL8, MET, CD40, LYN, IFI44, IFNGR1, EIF2AK2, IRF7, and IFIT5. These genes highlight pathways more focused on chemokine signaling, immune cell recruitment, and interferon-related responses.

The hub genes identified for Delta infections provide valuable insights into the severe disease manifestations commonly associated with this variant. Key genes such as TNFSF10 (TRAIL) and MYD88 suggest heightened apoptotic signaling and strong innate immune activation. TRAIL plays a role in inducing apoptosis in infected cells, which, while limiting viral spread, can also contribute to tissue damage, exacerbating disease severity (Zhu et al., 2019).

MYD88, a receptor for TLR8, a PRR for detection of viral mRNA and dsRNA in the cytoplasm early on in an infection, is crucial for mounting a rapid inflammatory response, but its overactivation can lead to cytokine storms, a hallmark of severe COVID-19 cases (Campbell et al., 2021; Kawai & Akira, 2010). Interferon-stimulated genes such as ISG15 and ISG20, further highlight robust antiviral defense mechanisms. However, prolonged activation of these pathways can lead to tissue damage and chronic inflammation, aligning with the extensive cytopathic effects observed in Delta infections (Tanneti et al., 2024). The inclusion of HLA-A, HLA-B, and PSMB9 highlights the importance of antigen presentation in Delta infections. These genes are pivotal for presenting viral antigens to T cells, facilitating an adaptive immune response.

However, excessive immune activation driven by these pathways may also result in immune-mediated damage. Transcription factors like IRF1 and NLRC5 are central to the regulation of these antiviral and inflammatory responses, underscoring Delta’s ability to drive both innate and adaptive immune responses, which may contribute to its more severe clinical manifestations compared to other variants (Choi & Smith, 2021).

In contrast, the hub genes identified for Omicron infections reflect its distinct pathogenic profile, characterized by high transmissibility but reduced disease severity in most cases.

Chemokines such as CXCL1 and CXCL8 (IL-8) play key roles in recruiting neutrophils and other immune cells to the site of infection, promoting robust immune surveillance in the early stages. MET and LYN, involved in cellular signaling and survival suggest mechanisms that support tissue repair and cellular homeostasis, potentially mitigating severe tissue damage observed in Delta infections (Hui et al., 2022). The interferon response remains critical in Omicron infections, as evidenced by the presence of IFI44, IFIT5, EIF2AK2, and IRF7. These genes are central to antiviral signaling and the orchestration of immune responses as they play key roles in detecting viral RNA (Jamilloux et al., 2020; Masood et al., 2021). CD40 and IFNGR1 reflect the activation of adaptive immune pathways, suggesting an efficient handoff between innate and adaptive immunity, potentially contributing to faster resolution of infection and reduced severity (McMahan et al., 2022). While Omicron’s ability to induce a robust interferon response early in the infection likely aids in controlling viral replication, this mechanism may also limit its pathogenicity in the lower respiratory tract.

Additionally, some of the identified hub genes play pivotal roles in the interferon (IFN) production pathway, as well as in key antiviral pathways such as the oligoadenylate-ribonuclease L (OAS-RNase L) activation pathway and the protein kinase R (PKR) activation pathway, underscoring their critical involvement in the host antiviral defense mechanisms (Y. Li et al., 2021). The IFN production pathway, involving hub genes such as ISG15, MYD88, ISG20, IL1R1, IRF1, and IRF7, leads to the secretion of antiviral interferon-stimulated genes (ISGs).

These ISGs mediate a wide array of antiviral activities, including viral RNA degradation, immune modulation, and the inhibition of viral replication. For instance, IRF1 and IRF7 are transcription factors that amplify IFN gene expression, ensuring a rapid and robust antiviral state, while MYD88 bridges Toll-like receptor signaling to IFN production, driving innate immune activation (Kawai & Akira, 2010). ISG15, in this pathway, functions as both a direct antiviral effector and a regulator of the innate immune response (Schoggins & Rice, 2011). The OAS-RNase L pathway, involving IFIT5 and IRF7, activates host RNase L, which degrades single-stranded RNA (ssRNA) indiscriminately. This mechanism not only targets viral RNA but can also limit host cell RNA to curtail viral replication. IFIT5, in particular, binds to viral RNA and enhances RNase L activity, while IRF7 facilitates upstream signaling that sustains this pathway’s activation (Y. Li et al., 2021). The PKR activation pathway, involving EIF2AK2, IFIT1, and ISG15, serves as a critical mechanism to block viral protein synthesis. Activation of EIF2AK2 (PKR) leads to phosphorylation of eIF2α, halting translation initiation and thereby inhibiting viral protein production. This pathway also induces stress granule formation, sequestering viral RNA and proteins to limit replication (Y. Li et al., 2021). IFIT1 and ISG15 further enhance this pathway’s antiviral effects by stabilizing RNA sensors and amplifying downstream signaling.

Overall, these interconnected pathways highlight the complexity of the antiviral response driven by these hub genes. The IFN production pathway establishes a broad antiviral state through ISGs, the OAS-RNase L pathway ensures degradation of viral RNA, and the PKR pathway halts viral translation, collectively creating a multilayered defense system. Their activation profiles may vary between variants like Delta and Omicron, contributing to differences in disease severity and immune evasion strategies.

## Limitations

The study relies on RNAseq data to analyze gene expression patterns and while this approach offers valuable insights into the transcriptome, it has inherent limitations. Firstly, identifying differential gene expression does not imply causation and so further wet lab validation experiments are required to establish cause-and-effect relationships and elucidate the functional consequences of these changes. The study has been conducted in vitro and while the ALI tissue culture captures the majority of gene expression observed in human airway, the model does not fully imitate an in vivo model. Gene co-expression analysis, performed to identify hub genes, also has specific limitations. While this approach identifies genes with correlated expression patterns, it does not necessarily confirm direct functional interactions or regulatory relationships. Furthermore, hub genes identified through co-expression analysis often reflect statistical centrality within the network rather than absolute biological importance, necessitating functional validation to confirm their roles. The choice of parameters for network construction, such as thresholds for correlation or module assignment, can also influence the resulting hub gene lists, introducing potential biases in interpretation.

## Conclusion

Our investigation uncovered notable differences in the impact of the Delta and Omicron variants of SARS-CoV-2 on host gene expression profiles. By utilizing differential gene expression (DEG) analysis and weighted gene co-expression network analysis (WGCNA) on RNA-seq data from air-liquid interface (ALI) cultures, we identified variant-specific hub genes and the pathways they regulate. Delta infections were characterized by a delayed immune response, with significant enrichment of pathways regulating cell death and apoptosis observed at 3 days post-infection (dpi). The hub genes identified were involved in apoptotic signaling, immune activation, and antigen presentation. These findings align with Delta’s clinical manifestations of severe disease, as these pathways regulate cytopathic effects, tissue damage, and heightened inflammation. Omicron infections showed an early induction of innate immune responses, with pathways related to interferon signaling, chemokine-mediated immune recruitment, and immune regulation activated as early as 24 hours post-infection. Hub genes highlighted Omicron’s robust interferon response and efficient immune activation, correlating with its high transmissibility and reduced lower respiratory tract involvement. The early immune activation likely limits Omicron’s pathogenicity compared to Delta, contributing to its association with milder disease outcomes despite its ability to evade some adaptive immune responses. By integrating transcriptomic data with gene network analysis, we have identified key hub genes and pathways that could serve as potential therapeutic targets or biomarkers for variant-specific interventions.

## Supplementary Figures

**Figure S1.**
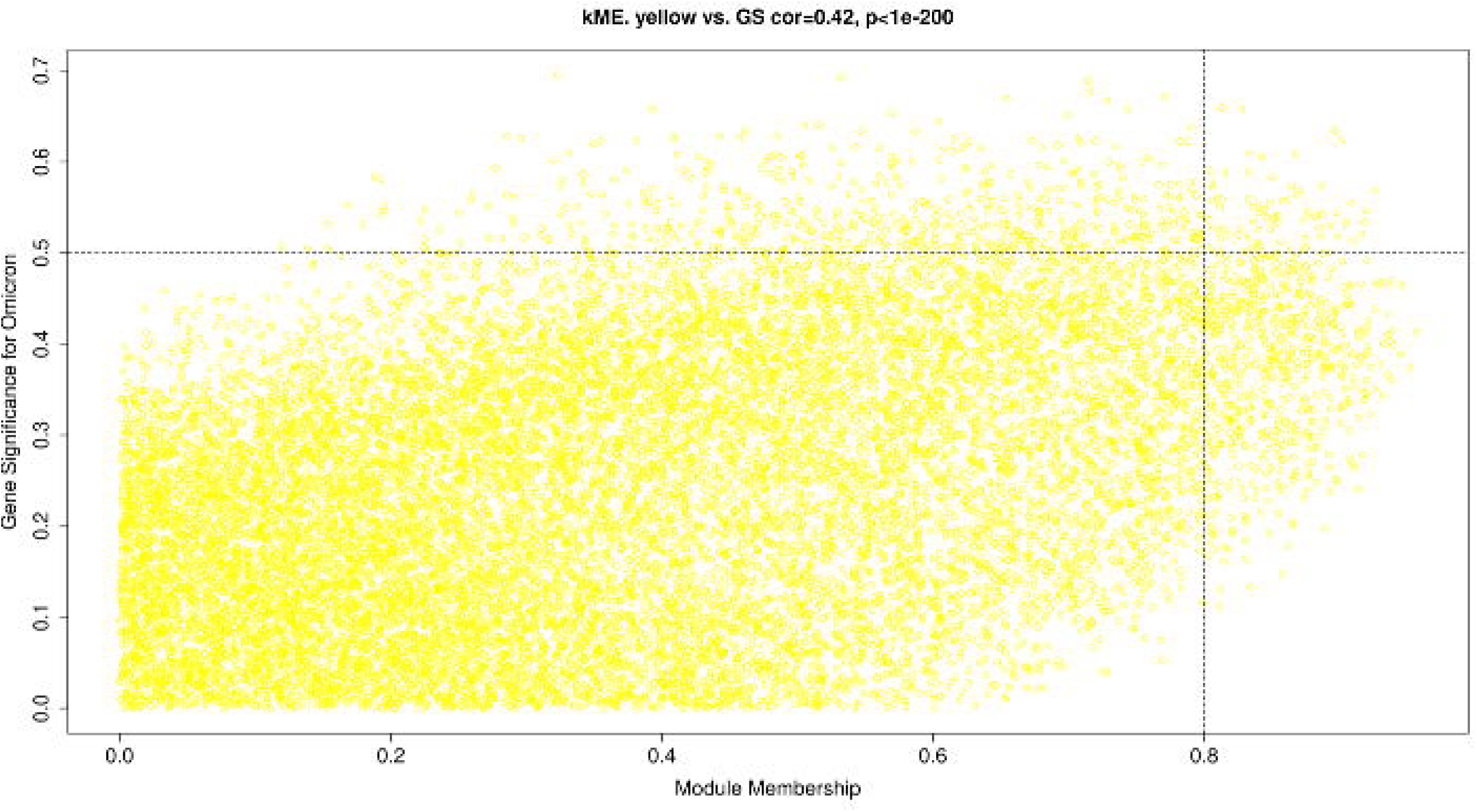
Volcano plot for fold change of human gene expression in ALI cultures infected with SARS-CoV-2 Delta and Omicron compared with healthy controls at 1, 2, and 3 dpi.

**Figure S2.**
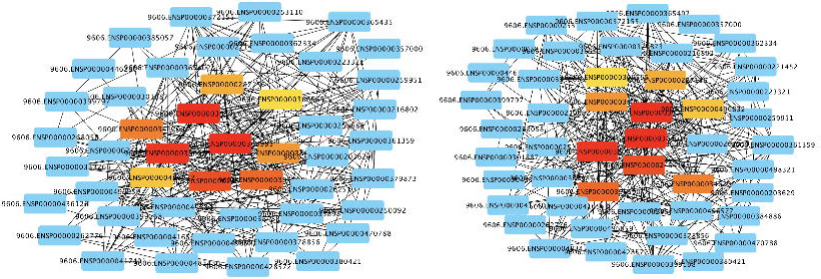
Identification of key modules correlated with SARS-CoV-2 Delta and Omicron infections using WGCNA. MM-GS plot for (a) Brown module, and (b) Yellow module.

**Figure S3.**
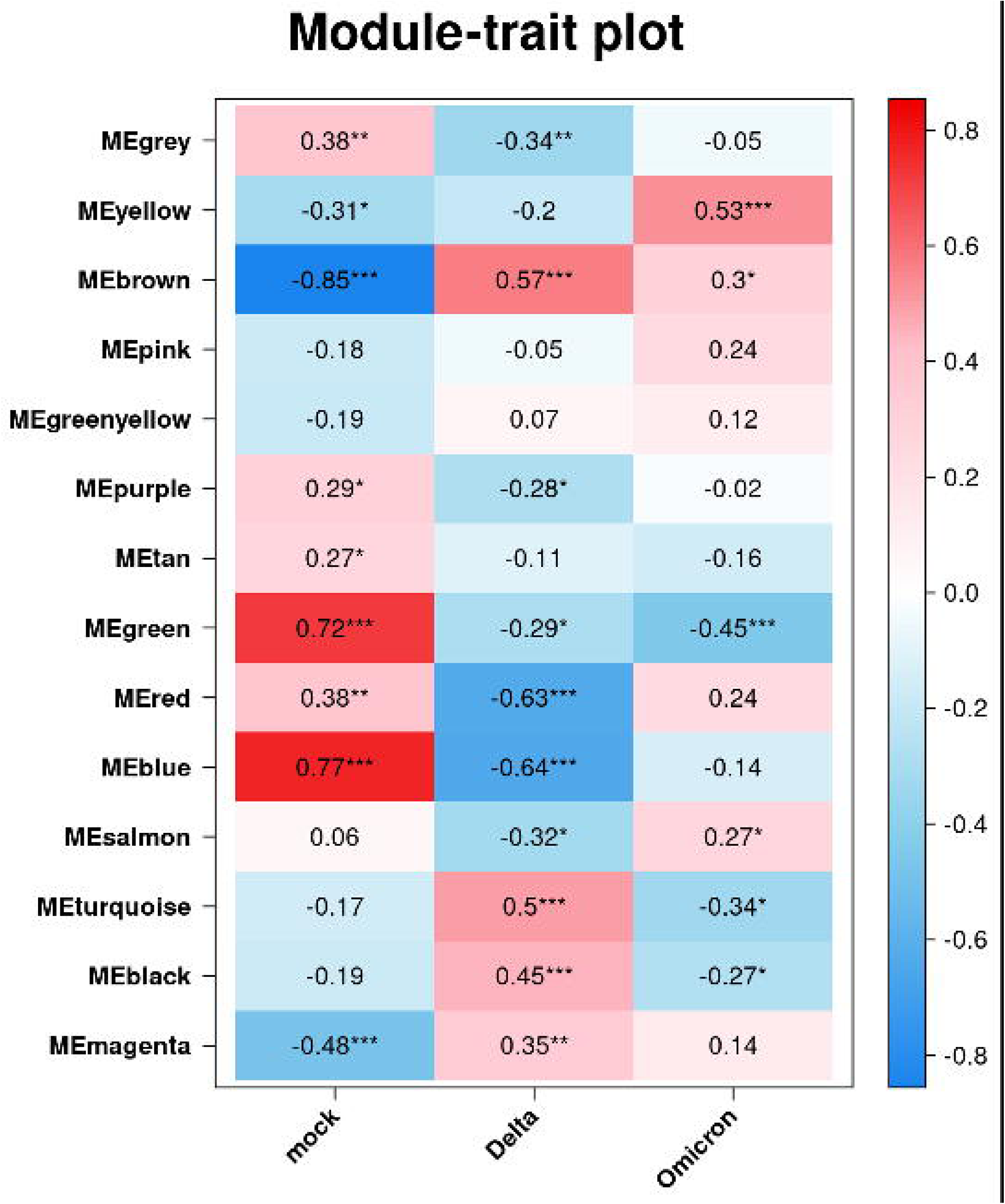
StringDB PPI network between candidate genes for (a) Brown module, and (b) Yellow module.

## References

1. Bajracharya, D., & Jansen, R. J. (2024). Observations of COVID-19 vaccine coverage and vaccine hesitancy on COVID-19 outbreak: An American ecological study. Vaccine, 42(2), 246–254. 10.1016/j.vaccine.2023.12.008

2. Bolger, A. M., Lohse, M., & Usadel, B. (2014). Trimmomatic: A flexible trimmer for Illumina sequence data. Bioinformatics, 30(15), 2114–2120. 10.1093/bioinformatics/btu170

3. Brüssow, H. (2022). COVID 19: Omicron – the latest, the least virulent, but probably not the last variant of concern of SARS CoV 2. Microbial Biotechnology, 15(7), 1927–1939. 10.1111/1751-7915.14064

4. Campbell, G. R., To, R. K., Hanna, J., & Spector, S. A. (2021). SARS-CoV-2, SARS-CoV-1, and HIV-1 derived ssRNA sequences activate the NLRP3 inflammasome in human macrophages through a non-classical pathway. iScience, *24*(4), 102295. 10.1016/j.isci.2021.102295

5. Castaneda, D. C., Jangra, S., Yurieva, M., Martinek, J., Callender, M., Coxe, M., Choi, A., García-Bernalt Diego, J., Lin, J., Wu, T.-C., Marches, F., Chaussabel, D., Yu, P., Salner, A., Aucello, G., Koff, J., Hudson, B., Church, S. E., Gorman, K., … Palucka, K. (2023). Spatiotemporally organized immunomodulatory response to SARS-CoV-2 virus in primary human broncho-alveolar epithelia. iScience, 26(8), 107374. 10.1016/j.isci.2023.107374

6. Chin, C.-H., Chen, S.-H., Wu, H.-H., Ho, C.-W., Ko, M.-T., & Lin, C.-Y. (2014). cytoHubba: Identifying hub objects and sub-networks from complex interactome. BMC Systems Biology, 8(S4), S11. 10.1186/1752-0509-8-S4-S11

7. Choi, J. Y., & Smith, D. M. (2021). SARS-CoV-2 Variants of Concern. Yonsei Medical Journal, 62(11), 961. 10.3349/ymj.2021.62.11.961

8. Di Tommaso, P., Chatzou, M., Floden, E. W., Barja, P. P., Palumbo, E., & Notredame, C. (2017). Nextflow enables reproducible computational workflows. Nature Biotechnology, 35(4), 316–319. 10.1038/nbt.3820

9. Ghosh, B., Park, B., Bhowmik, D., Nishida, K., Lauver, M., Putcha, N., Gao, P., Ramanathan, M., Hansel, N., Biswal, S., & Sidhaye, V. K. (2020). Strong correlation between air-liquid interface cultures and in vivo transcriptomics of nasal brush biopsy. American Journal of Physiology-Lung Cellular and Molecular Physiology, 318(5), L1056–L1062. 10.1152/ajplung.00050.2020

10. Girona-Alarcon, M., Argüello, G., Esteve-Sole, A., Bobillo-Perez, S., Burgos-Artizzu, X. P., Bonet-Carne, E., Mensa-Vilaró, A., Codina, A., Hernández-Garcia, M., Jou, C., Alsina, L., & Jordan, I. (2022). Low levels of CIITA and high levels of SOCS1 predict COVID- 19 disease severity in children and adults. iScience, 25(1), 103595. 10.1016/j.isci.2021.103595

11. He, X., Hong, W., Pan, X., Lu, G., & Wei, X. (2021). SARS CoV 2 Omicron variant: Characteristics and prevention. MedComm, 2(4), 838–845. 10.1002/mco2.110

12. Hui, K. P. Y., Ho, J. C. W., Cheung, M., Ng, K., Ching, R. H. H., Lai, K., Kam, T. T., Gu, H., Sit, K.-Y., Hsin, M. K. Y., Au, T. W. K., Poon, L. L. M., Peiris, M., Nicholls, J. M., & Chan, M. C. W. (2022). SARS-CoV-2 Omicron variant replication in human bronchus and lung ex vivo. Nature, 603(7902), 715–720. 10.1038/s41586-022-04479-6

13. Jamilloux, Y., Henry, T., Belot, A., Viel, S., Fauter, M., El Jammal, T., Walzer, T., François, B., & Sève, P. (2020). Should we stimulate or suppress immune responses in COVID-19? Cytokine and anti-cytokine interventions. Autoimmunity Reviews, 19(7), 102567. 10.1016/j.autrev.2020.102567

14. Karami, H., Derakhshani, A., Ghasemigol, M., Fereidouni, M., Miri-Moghaddam, E., Baradaran, B., Tabrizi, N., Najafi, S., Solimando, A., Marsh, L., Silvestris, N., De Summa, S., Paradiso, A., Racanelli, V., & Safarpour, H. (2021). Weighted Gene Co-Expression Network Analysis Combined with Machine Learning Validation to Identify Key Modules and Hub Genes Associated with SARS-CoV-2 Infection. Journal of Clinical Medicine, 10(16), 3567. 10.3390/jcm10163567

15. Kawai, T., & Akira, S. (2010). The role of pattern-recognition receptors in innate immunity: Update on Toll-like receptors. Nature Immunology, 11(5), 373–384. 10.1038/ni.1863

16. Kim, D., Paggi, J. M., Park, C., Bennett, C., & Salzberg, S. L. (2019). Graph-based genome alignment and genotyping with HISAT2 and HISAT-genotype. Nature Biotechnology, 37(8), 907–915. 10.1038/s41587-019-0201-4

17. Langfelder, P., & Horvath, S. (2008). WGCNA: An R package for weighted correlation network analysis. BMC Bioinformatics, 9(1), 559. 10.1186/1471-2105-9-559

18. Lee, K. S., Wong, T. Y., Russ, B. P., Horspool, A. M., Miller, O. A., Rader, N. A., Givi, J. P., Winters, M. T., Wong, Z. Y. A., Cyphert, H. A., Denvir, J., Stoilov, P., Barbier, M., Roan, N. R., Amin, Md. S., Martinez, I., Bevere, J. R., & Damron, F. H. (2022). SARS-CoV-2 Delta variant induces enhanced pathology and inflammatory responses in K18-hACE2 mice. PLOS ONE, 17(8), e0273430. 10.1371/journal.pone.0273430

19. Li, H., Handsaker, B., Wysoker, A., Fennell, T., Ruan, J., Homer, N., Marth, G., Abecasis, G., Durbin, R., & 1000 Genome Project Data Processing Subgroup. (2009). The Sequence Alignment/Map format and SAMtools. Bioinformatics, *25*(16), 2078–2079. 10.1093/bioinformatics/btp352

20. Li, Y., Renner, D. M., Comar, C. E., Whelan, J. N., Reyes, H. M., Cardenas-Diaz, F. L., Truitt, R., Tan, L. H., Dong, B., Alysandratos, K. D., Huang, J., Palmer, J. N., Adappa, N. D., Kohanski, M. A., Kotton, D. N., Silverman, R. H., Yang, W., Morrisey, E. E., Cohen, N. A., & Weiss, S. R. (2021). SARS-CoV-2 induces double-stranded RNA-mediated innate immune responses in respiratory epithelial-derived cells and cardiomyocytes. Proceedings of the National Academy of Sciences, 118(16), e2022643118. 10.1073/pnas.2022643118

21. Liberzon, A., Birger, C., Thorvaldsdóttir, H., Ghandi, M., Mesirov, J. P., & Tamayo, P. (2015). The Molecular Signatures Database Hallmark Gene Set Collection. Cell Systems, 1(6), 417–425. 10.1016/j.cels.2015.12.004

22. Love, M. I., Huber, W., & Anders, S. (2014). Moderated estimation of fold change and dispersion for RNA-seq data with DESeq2. Genome Biology, 15(12), 550. 10.1186/s13059-014-0550-8

23. Masood, K. I., Yameen, M., Ashraf, J., Shahid, S., Mahmood, S. F., Nasir, A., Nasir, N., Jamil, B., Ghanchi, N. K., Khanum, I., Razzak, S. A., Kanji, A., Hussain, R., E. Rottenberg, M., & Hasan, Z. (2021). Upregulated type I interferon responses in asymptomatic COVID-19 infection are associated with improved clinical outcome. Scientific Reports, 11(1), 22958. 10.1038/s41598-021-02489-4

24. McMahan, K., Giffin, V., Tostanoski, L. H., Chung, B., Siamatu, M., Suthar, M. S., Halfmann, P., Kawaoka, Y., Piedra-Mora, C., Jain, N., Ducat, S., Kar, S., Andersen, H., Lewis, M. G., Martinot, A. J., & Barouch, D. H. (2022). Reduced pathogenicity of the SARS-CoV-2 omicron variant in hamsters. Med, 3(4), 262–268.e4. 10.1016/j.medj.2022.03.004

25. Naqvi, A. A., Fatima, K., Mohammad, T., Fatima, U., Singh, I. K., Singh, A., Atif, S. M., Hariprasad, G., Hasan, G. M., & Hassan, Md. I. (2020). Insights into SARS-COV-2 genome, structure, evolution, pathogenesis and therapies: Structural Genomics Approach. Biochimica et Biophysica Acta (BBA) - Molecular Basis of Disease, 1866(10), 165878. 10.1016/j.bbadis.2020.165878

26. Pertea, M., Pertea, G. M., Antonescu, C. M., Chang, T.-C., Mendell, J. T., & Salzberg, S. L. (2015). StringTie enables improved reconstruction of a transcriptome from RNA-seq reads. Nature Biotechnology, 33(3), 290–295. 10.1038/nbt.3122

27. Rayner, R. E., Makena, P., Prasad, G. L., & Cormet-Boyaka, E. (2019). Optimization of Normal Human Bronchial Epithelial (NHBE) Cell 3D Cultures for in vitro Lung Model Studies. Scientific Reports, 9(1), 500. 10.1038/s41598-018-36735-z

28. Saichi, M., Ladjemi, M. Z., Korniotis, S., Rousseau, C., Ait Hamou, Z., Massenet-Regad, L., Amblard, E., Noel, F., Marie, Y., Bouteiller, D., Medvedovic, J., Pène, F., & Soumelis, V. (2021). Single-cell RNA sequencing of blood antigen-presenting cells in severe COVID-19 reveals multi-process defects in antiviral immunity. Nature Cell Biology, 23(5), 538–551. 10.1038/s41556-021-00681-2

29. Sayers, E. W., Bolton, E. E., Brister, J. R., Canese, K., Chan, J., Comeau, D. C., Connor, R., Funk, K., Kelly, C., Kim, S., Madej, T., Marchler-Bauer, A., Lanczycki, C., Lathrop, S., Lu, Z., Thibaud-Nissen, F., Murphy, T., Phan, L., Skripchenko, Y., … Sherry, S. T. (2022). Database resources of the national center for biotechnology information. Nucleic Acids Research, 50(D1), D20–D26. 10.1093/nar/gkab1112

30. Schoggins, J. W., & Rice, C. M. (2011). Interferon-stimulated genes and their antiviral effector functions. Current Opinion in Virology, 1(6), 519–525. 10.1016/j.coviro.2011.10.008

31. Shannon, P., Markiel, A., Ozier, O., Baliga, N. S., Wang, J. T., Ramage, D., Amin, N., Schwikowski, B., & Ideker, T. (2003). Cytoscape: A Software Environment for Integrated Models of Biomolecular Interaction Networks. Genome Research, 13(11), 2498–2504. 10.1101/gr.1239303

32. Subramanian, A., Tamayo, P., Mootha, V. K., Mukherjee, S., Ebert, B. L., Gillette, M. A., Paulovich, A., Pomeroy, S. L., Golub, T. R., Lander, E. S., & Mesirov, J. P. (2005). Gene set enrichment analysis: A knowledge-based approach for interpreting genome-wide expression profiles. Proceedings of the National Academy of Sciences, 102(43), 15545– 15550. 10.1073/pnas.0506580102

33. Szklarczyk, D., Gable, A. L., Nastou, K. C., Lyon, D., Kirsch, R., Pyysalo, S., Doncheva, N. T., Legeay, M., Fang, T., Bork, P., Jensen, L. J., & von Mering, C. (2021). The STRING database in 2021: Customizable protein–protein networks, and functional characterization of user-uploaded gene/measurement sets. Nucleic Acids Research, 49(D1), D605–D612. 10.1093/nar/gkaa1074

34. Tanneti, N. S., Patel, A. K., Tan, L. H., Marques, A. D., Perera, R. A. P. M., Sherrill-Mix, S., Kelly, B. J., Renner, D. M., Collman, R. G., Rodino, K., Lee, C., Bushman, F. D., Cohen, N. A., & Weiss, S. R. (2024). Comparison of SARS-CoV-2 variants of concern in primary human nasal cultures demonstrates Delta as most cytopathic and Omicron as fastest replicating. mBio, 15(4), e03129–23. 10.1128/mbio.03129-23

35. Thaker, S. K., Ch’ng, J., & Christofk, H. R. (2019). Viral hijacking of cellular metabolism. BMC Biology, 17(1), 59. 10.1186/s12915-019-0678-9

36. Ulloa, A. C., Buchan, S. A., Daneman, N., & Brown, K. A. (2022). Estimates of SARS-CoV-2 Omicron Variant Severity in Ontario, Canada. JAMA, 327(13), 1286. 10.1001/jama.2022.2274

37. Goethem, N V., Chung, P. Y. J., Meurisse, M., Vandromme, M., De Mot, L., Brondeel, R., Stouten, V., Klamer, S., Cuypers, L., Braeye, T., Catteau, L., Nevejan, L., Van Loenhout, J. A. F., & Blot, K. (2022). Clinical Severity of SARS-CoV-2 Omicron Variant Compared with Delta among Hospitalized COVID-19 Patients in Belgium during Autumn and Winter Season 2021–2022. Viruses, 14(6), 1297. 10.3390/v14061297

38. WHO Coronavirus (COVID-19) Dashboard | WHO Coronavirus (COVID-19) Dashboard With Vaccination Data. (n.d.). Retrieved February 20, 2022, from https://covid19.who.int/

39. Wu, T., Hu, E., Xu, S., Chen, M., Guo, P., Dai, Z., Feng, T., Zhou, L., Tang, W., Zhan, L., Fu, X., Liu, S., Bo, X., & Yu, G. (2021). clusterProfiler 4.0: A universal enrichment tool for interpreting omics data. The Innovation, 2(3), 100141. 10.1016/j.xinn.2021.100141

40. Zhu, Z.-C., Liu, J.-W., Yang, C., Li, M.-J., Wu, R.-J., & Xiong, Z.-Q. (2019). Targeting KPNB1 overcomes TRAIL resistance by regulating DR5, Mcl-1 and FLIP in glioblastoma cells. Cell Death & Disease, *10*(2), 118. 10.1038/s41419-019-1383-x

